# Feeding desensitizes A1 adenosine receptors in adipose through FOXO1-mediated transcriptional regulation

**DOI:** 10.1101/2022.03.04.482364

**Authors:** Mitchell E. Granade, Stefan R. Hargett, Michael C. Lemke, Melissa A. Luse, Irina M. Bochkis, Joel Linden, Thurl E. Harris

## Abstract

Adipose tissue is a critical regulator of energy balance that must rapidly shift its metabolism between fasting and feeding to maintain homeostasis. Adenosine has been characterized as an important regulator of adipocyte metabolism primarily through its actions on A_1_ adenosine receptors (A1R). We sought to understand the role A1R plays in adipocytes during fasting and feeding to regulate glucose and lipid metabolism by using an inducible, adiponectin-Cre with *Adora1* floxed mice (FAdora1^−/−^), where F designates a fat-specific deletion. Fadora1^−/−^ mice had impairments in the suppression of lipolysis by insulin on normal chow and impaired glucose tolerance on high-fat diet. FAdora1^−/−^ mice also exhibited a higher lipolytic response to isoproterenol than WT controls when fasted, but not after a 4-hour refeeding period. We found that FOXO1 binds to the A1R promoter in adipocytes. Upon feeding, signaling along the insulin-Akt-FOXO1 axis leads to a rapid downregulation of A1R transcript and desensitization of adipocytes to A1R agonism. Obesity also desensitizes adipocyte A1R, and this is accompanied by a disruption of cyclical changes in A1R transcription between fasting and refeeding. We propose that FOXO1 drives high A1R expression under fasted conditions to limit excess lipolysis during stress and augment insulin action upon feeding. Subsequent downregulation of A1R under fed conditions facilitates reentrance into the catabolic state upon fasting.

## Introduction

Adipose tissue is a critical regulator of energy balance, and must rapidly adapt its metabolic function to either esterify fatty acids for adequate energy storage during feeding or undergo lipolysis to provide non-esterified fatty acids (NEFA) and glycerol for energy during fasting (1). These homeostatic set points are primarily regulated by catabolic hormones, such as catecholamines, driving lipolysis, and anabolic hormones, such as insulin, suppressing lipolysis and driving fatty acid uptake and esterification (2).

Adenosine is a nucleoside with a role in extracellular signaling that has been well established as an important regulator of adipocyte function through opposing catecholamine signaling during stress and enhancing insulin signaling during feeding (3). Adenosine signals through four G protein-coupled receptors: A_1_, A_2A_, A_2B_, and A_3_ (4). In adipocytes, adenosine has been found to act primarily through the A_1_ adenosine receptor (A1R), a Gαi-coupled receptor that acts to reduce cyclic AMP (cAMP), suppress lipolysis and counteract ß-adrenergic signaling (5). Adenosine signaling via A1R in adipocytes has been shown to augment insulin stimulated glucose uptake, lipogenesis, and suppression of lipolysis *in vitro* (6–8). Agonism of A1R has also been shown to reduce serum NEFA in both rats and humans, and improve insulin sensitivity in obese rats (9–11). Further, reduced A1R expression in adipocytes due to obesity, hypertension, or prolonged pharmacological stimulation are associated with insulin resistance (5,12,13). Despite this, mice with global A1R knock out display relatively normal systemic NEFA levels, and differences in glucose and insulin tolerance were concomitant with significant confounding factors such as differences in body weight, food intake, insulin and glucagon secretion, and survival (6–10). It therefore remains unclear whether endogenous adenosine and A1R contribute to shifts in adipocyte metabolism between the fasted and fed state *in vivo*.

Adipose tissue maintains basal levels of adenosine that are considered sufficient to prevent a maximal stimulation of lipolysis through tonic activation of A1R (11–15). However, adenosine has long been associated with stress responses and adipose interstitial levels of adenosine increase drastically in response to adrenergic stress (16). In adipocytes, the degradation of adenosine with exogenous adenosine deaminase or inhibition with an A1R antagonist enhances the stimulation of lipolysis by adrenergic signals such as norepinephrine or the β-adrenergic receptor agonist isoproterenol (13–15,17). This suggests that A1R activation may serve as a protective response since high levels of NEFA resulting from stress-induced lipolysis can have deleterious effects such as insulin resistance and lipotoxicity (18,19). However, the effects of A1R signaling in adipocytes in response to endogenous adenosine and subsequent impacts to circulating NEFA during stress have not been investigated.

We sought to explore how the loss of A1R specifically in adipose tissue affects glucose metabolism in lean and obese mice. We also sought to understand how the loss of adipose A1R affects lipid metabolism in the fed and fasted states, as well as the lipolytic response to stress. In this study, we show that A1R acts to augment insulin action to suppress lipolysis in the transition from the fasted to the fed state, and loss of adipose A1R leads to glucose intolerance in obese mice. We also demonstrate that A1R limits the lipolytic response to stress in the fasted, but not fed, state. We identified a mechanism of differential regulation of A1R during the fasted and fed states whereby insulin downregulates A1R in adipocytes and desensitizes adipose to A1R agonism in the fed state.

## Research Design and Methods

### Animals

All animals were bred and maintained in accordance with the University of Virginia Animal Care and Use Committee regulations and the study was approved by the ACUC ethics committee. 12- to 16-week-old male and female mice were used for studies unless otherwise indicated. *Adora1*^loxP/loxP^ mice were generously shared with us by Dr. Stanislav Zakharenko (20). Inducible, fat-specific *Adora1* knockout mice were generated by crossing *Adora1*^loxP/loxP^ with AdipoQ-Cre-ERT2 mice resulting in wildtype (WT) *Adora1*^loxP/loxP^ mice and knockout *Adora1*^loxP/loxP^;AdipoQ-Cre-ERT2 (FAdora1^−/−^) littermates on a mixed C57BL/6J and C57BL/6N background.

### In vivo measurements

Heart rate was measured by electrocardiogram in anesthetized mice. Fat and lean mass distribution were determined using Echo-MRI. Ambulatory activity, carbon dioxide production (VCO_2_), and oxygen consumption (VO_2_) were measured using an Oxymax metabolic chamber system.

### Measurement of glucose, insulin, and NEFA

Blood glucose was measured using a One-Touch Ultra glucometer. Insulin and NEFA levels were measured in serum samples or lipolysis assay buffer using the Ultra-Sensitive Mouse Insulin ELISA Kit (Crystal Chem) and the HR series NEFA-HR(2) Kit (Wako Diagnostics) respectively and according to the manufacturer’s protocols.

### RNA extraction and real-time qPCR

Total RNA was isolated using the PureLink RNA Mini Kit (Invitrogen) according to the manufacturer’s protocol. For tissues, RNA was first extracted using Trizol Reagent (Invitrogen). cDNA was synthesized from 500 ng RNA with the High-Capacity RNA-to-cDNA Synthesis Kit (Applied Biosystems). Real-time qPCR was performed with the iQ SYBR Green Supermix (Bio-Rad) in duplicate and analyzed with a CFX96 Real-Time PCR Detection System (Bio-Rad). Data were normalized to *Ppia* cDNA using the delta-delta-Ct method.

### Western blot and quantitative analysis

Equal amounts of proteins were separated by SDS-PAGE and transferred to PVDF. Membranes were blocked with 10% dry milk in TBST, and western blotting was carried out with appropriate primary and secondary antibodies. Blots were imaged by detecting horseradish peroxidase conjugated chemiluminescence on an Amersham ImageQuant 800. For assessing differences in protein or phospho-protein signal, membranes were stripped and reprobed with β-tubulin, α-actin, or TBP loading controls as appropriate and as indicated, and phospho-protein blots were additionally reprobed with the corresponding total antibody. Blots were quantified using densitometry with ImageQuant software.

### Chromatin immunoprecipitation

Chromatin immunoprecipitation (ChIP) was performed as previously described with modifications (21). Briefly, 3T3-L1 adipocytes were serum starved overnight in DMEM with 0.25% FBS and 0.25% BSA and treated with vehicle or 1 μmol/L AS-1842856 for 4 hours. Cells were fixed in 1% formaldehyde for 10 minutes, and fixation was quenched with 125 mmol/L glycine for 5 minutes. Cells were washed twice with cold PBS, lysed in hypotonic lysis buffer (10 mmol/L Tris-HCl, pH 8.0, 10 mmol/L NaCl, 3 mmol/L MgCl_2_, 0.5% NP-40) supplemented with protease inhibitors by Dounce homogenization, and incubated on ice for 5 minutes. Nuclei were pelleted by centrifugation at 17,000 × *g* for 5 minutes at 4°C. Pellets were resuspended with SDS lysis buffer (50 mmol/L Tris-HCl, pH 8.0, 10 mmol/L EDTA, 1% SDS) and chromatin was sheared by sonication using a Diagenode BioRuptor Pico with 30 sec on and 30 sec off for 12 cycles. ChIP was performed using 3 μg of FOXO1 rabbit-antibody (ProteinTech) or normal rabbit IgG (Cell Signaling), and antibodies were pulled down with protein A agarose (Roche). Chromatin pulldown of the *Adora1* promoter region or 28S rDNA control region was assessed by qPCR using the following primers: *Adora1* promoter: 5’-TGACCCTTGAAACCATGTGA-3’ and 5’-CCCAGAGTACCCAACACACA-3’, 28S rDNA: 5’-CTGGGTATAGGGGCGAAAGAC-3’ and 5’-GGCCCCAAGACCTCTAATCAT-3’. FOXO1 binding was measured as enrichment over normal rabbit IgG ChIP.

## Results

### Loss of A1R inhibition of lipolysis in adipocyte-specific A1R knockout mice

Several groups have previously evaluated the role of A1R in adipose tissue and fatty acid metabolism using mice with a global knockout of A1R, however these mice exhibit increased body weight and food intake (8,9), increased insulin and glucagon secretion (7,10), and decreased survival (6,22) which make it difficult to interpret the role of A1R specifically in adipose. To avoid these complications, we crossed mice with a tamoxifen-inducible Cre driven by the adiponectin promoter (Jax #025124) with an *Adora1*^loxP*/*loxP^ strain (20) in which the terminal 3’ exon of the *Adora1* gene, which encodes A1R, is flanked by loxP sites. Both Cre negative *Adora1*^loxP*/*loxP^ control mice (WT) and Cre positive *Adora1*^loxP*/*loxP^;AdipoQ-Cre-ERT2 knockout mice (FAdora1^−/−^), where F indicates a fat-specific deletion, were treated with tamoxifen for 10 days at 8-weeks of age to induce knockout, followed by a 2-week washout period. Following tamoxifen treatment, FAdora1^−/−^ mice exhibited an ~88% knockdown in A1R mRNA in isolated white adipocytes from gonadal fat pads and a ~95% knockdown in brown adipose tissue (Fig. 1A and B). This resulted in an ~80% reduction in protein levels within isolated white adipocytes (Fig. 1C). The FAdora1^−/−^ mice did not display any compensatory changes in the other adenosine receptors A_2A_, A_2B_, or A_3_ in white adipocytes or brown adipose tissue (Fig. S1). Meanwhile, A1R is also highly expressed in cardiac tissue and no reduction in A1R protein levels was observed within whole-heart lysates (Fig. 1D) (23).

**Figure 1:**
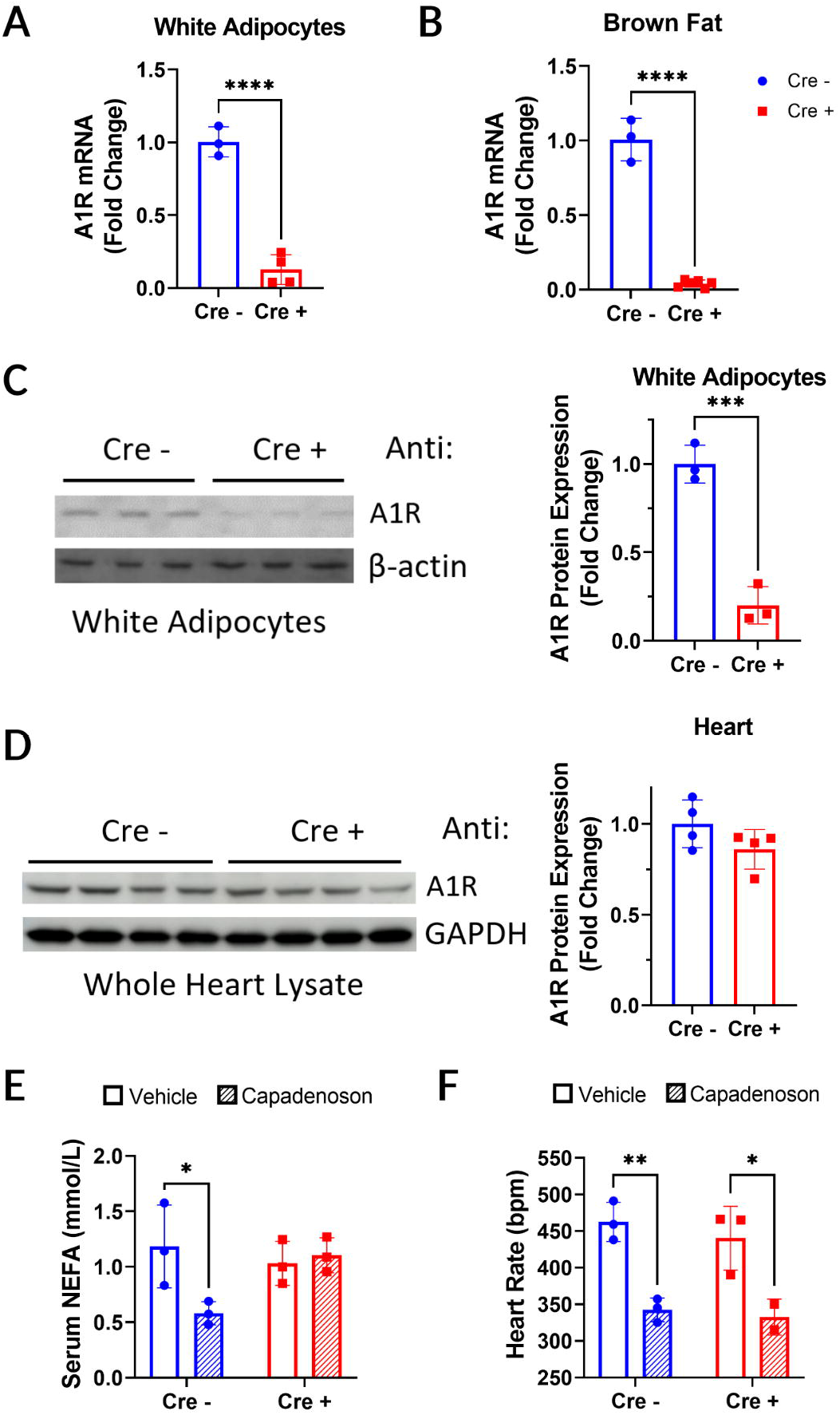
Tamoxifen induces a functional knockout of adipose A1R in FAdora1^−/−^ mice. (A and B) mRNA levels of A1R in Cre negative WT (blue) and Cre positive FAdora1^−/−^ mice (red) in isolated epididymal white adipocytes (A) and brown adipose tissue (B). (C) Protein levels of A1R in isolated gonadal white adipocytes from WT and FAdora1^−/−^ mice assessed and quantified by western blot normalized to β-actin loading control. (d) Protein levels of A1R in whole heart lysates from WT and FAdora1^−/−^ mice assessed and quantified by western blot normalized to GAPDH loading control. (E and F) Serum NEFA levels (E) and heart rate (F) in WT and FAdora1^−/−^ mice fasted overnight and treated for 2 hours with vehicle or 1 mg/kg capadenoson. Error bars represent SD. *p<0.05, **p<0.01, ***p<0.001, ****p<0.0001, t-test or two-way ANOVA.

Agonism of A1R in adipocytes results in lower levels of non-esterified fatty acids (NEFA) within the blood stream (24). Partial agonists of A1R such as capadenoson have previously been shown to reduce serum NEFA levels as well as cause modest reductions in heart rate through A1 receptors in the AV node (23,25). We subjected WT and FAdora1^−/−^ mice to an overnight fast and treated with either vehicle or capadenoson for 2 hours. Capadenoson significantly reduced serum NEFA in WT mice, but not FAdora1^−/−^ mice (Fig. 1E). Consistent with adipocyte-specific loss of A1R in our system, capadenoson treatment reduced heart rate in both WT and FAdora1^−/−^ mice (Fig. 1F).

### Impaired insulin suppression of lipolysis in male FAdora1^−/−^ mice

In contrast to global A1R knockout mice, FAdora1^−/−^ mice did not develop changes in body weight or fat and lean mass distribution following tamoxifen administration, even in a long-term follow-up more than 30 weeks after A1R knockout (Fig. 2A and B, Fig. S2). There were no differences between WT and FAdora1^−/−^ male mice in blood glucose or serum insulin when fasted overnight, and only minor increases in glucose and insulin levels following a 4-hour refeeding period (Fig. 2C and D). There were also no differences in fasted or refed NEFA levels (Fig. 2E), glucose (Fig. 2F) or insulin secretion (Fig. 2G) during a glucose tolerance test (GTT), or insulin tolerance (Fig. 2H). Overall, FAdora1^−/−^ mice were relatively metabolically normal. However, when we treated fasted WT and FAdora1^−/−^ male mice with insulin for 15 minutes, serum NEFA was significantly reduced in WT but not FAdora1^−/−^ mice (Fig 2I). This suggests that the loss of A1R in adipocytes impairs insulin action to suppress lipolysis. Throughout our characterization of the FAdora1^−/−^ mice, female mice consistently lacked a distinct phenotype as compared with male mice. Female FAdora1^−/−^ mice did not display significant differences in body weight, body mass distribution, fasting and refed values for glucose, insulin and NEFA, glucose tolerance, or insulin secretion (Fig. S3).

**Figure 2:**
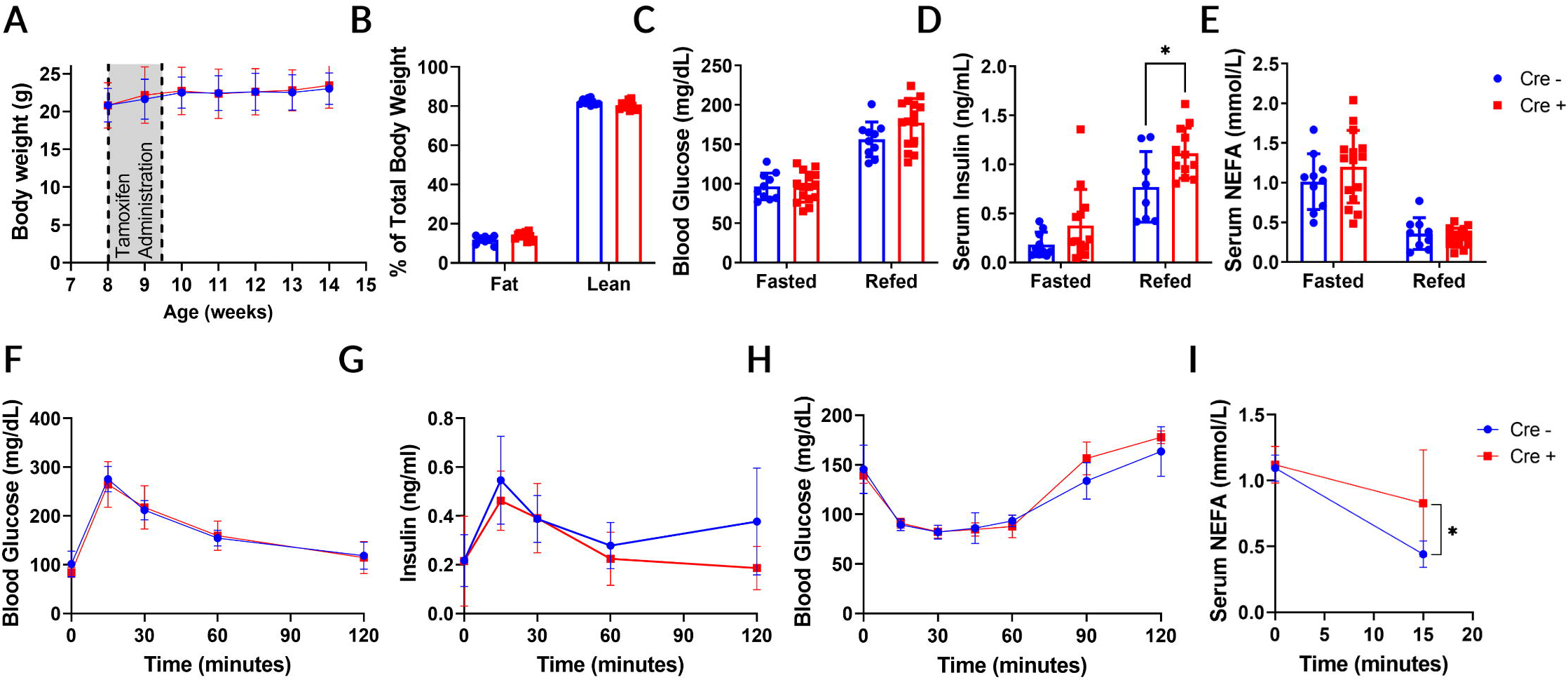
Induced loss of A1R in adipose impairs insulin suppression of lipolysis. (A) Body weight of male Cre negative WT (blue) and Cre positive FAdora1^−/−^ mice (red) starting at the time of tamoxifen administration at 8 weeks. Gray box indicates timing of tamoxifen. (B) Fat and lean mass distribution in male WT and FAdora1^−/−^ mice 6 weeks after the start of tamoxifen administration measured by EchoMRI. (C-E) Blood glucose (C), serum insulin (D), and serum NEFA (E) levels in male WT and FAdora1^−/−^ mice after an overnight fast (fasted) and 4-hour refeeding period (refed). (F, G) Blood glucose (F) and serum insulin (G) in male WT and FAdora1^−/−^ mice during an i.p. glucose tolerance test with 1 g/kg glucose. (H) Blood glucose in male WT and FAdora1^−/−^ mice during an i.p. insulin tolerance test with 0.75 IU/kg insulin. (I) Serum NEFA in overnight fasted male WT and FAdora1^−/−^ mice at 0 and 15 minutes after administration of 0.75 IU/mL insulin. Error bars represent SD. *p<0.05, t-test or two-way ANOVA.

### High fat diet results in overt glucose intolerance within male FAdora1^−/−^ mice

To further explore the impacts of A1R knockout on adipose insulin sensitivity, we placed WT and FAdora1^−/−^ mice on high fat diet (HFD) for 12 weeks. Tamoxifen was administered beginning after 7 weeks of HFD feeding to allow sufficient time for tamoxifen washout and recovery. While tamoxifen had a clear secondary impact on weight gain, there were no differences between WT and FAdora1^−/−^ male mice in body weight or body mass distribution (Fig. 3A and B). In addition, FAdora1^−/−^ male mice did not have any differences in activity or respiratory exchange ratio (Fig. 3C and D). There also continued to be no differences in fasting or refed glucose, insulin, and NEFA levels (Fig. 3E–G). However, while there were no differences in insulin tolerance (Fig. 3H), HFD fed FAdora1^−/−^ mice displayed clear glucose intolerance during an i.p. GTT with no change in insulin secretion (Fig. 3I and J). There were no significant differences in metabolic parameters between WT and FAdora1^−/−^ among HFD fed female mice (Fig. S4).

**Figure 3:**
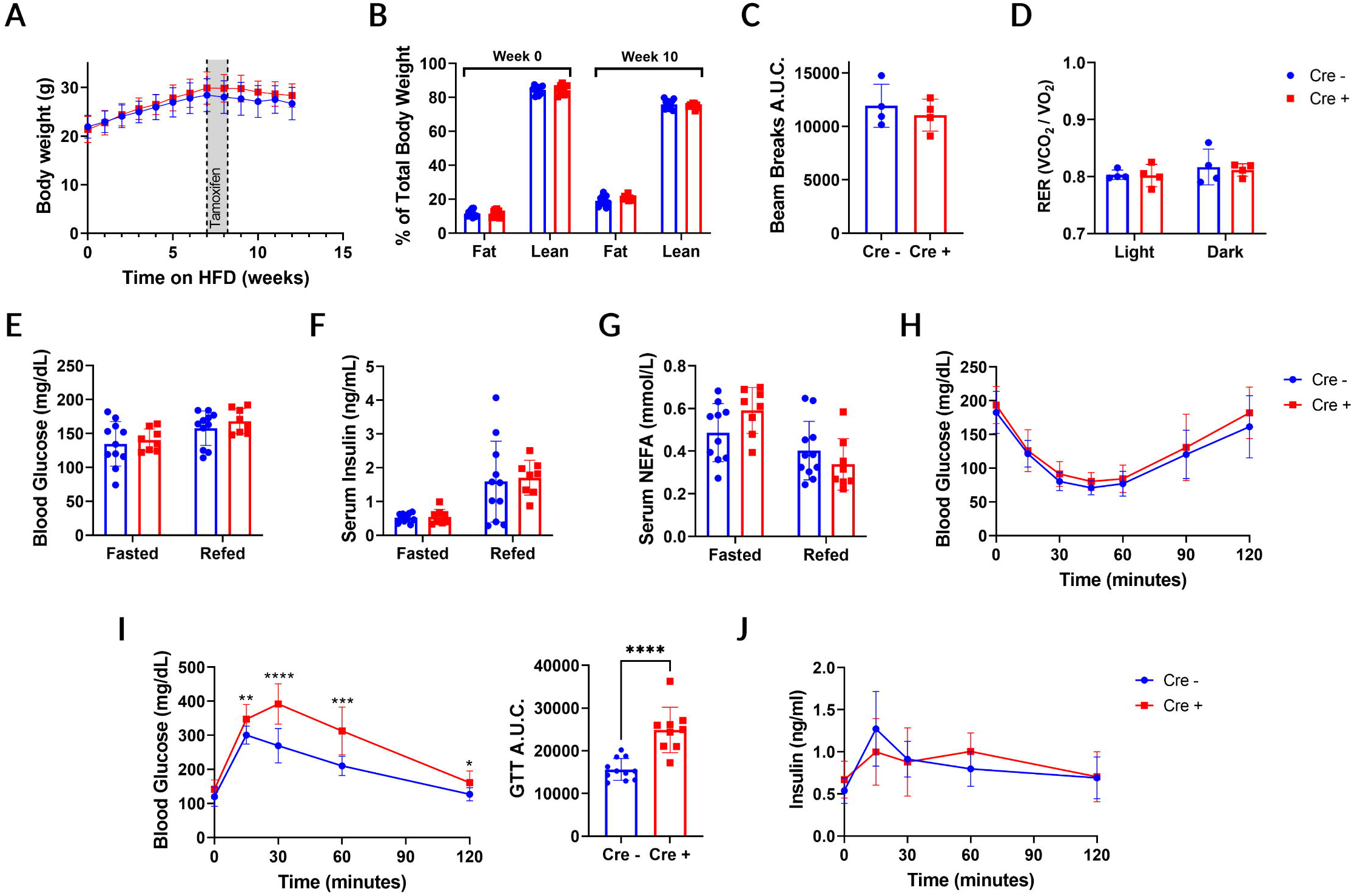
FAdora1^−/−^ mice are more susceptible to high-fat diet-induced glucose intolerance. (A) Body weight of male Cre negative WT (blue) and Cre positive FAdora1^−/−^ mice (red) starting at the beginning of high-fat diet feeding at 8 weeks. Gray box indicates timing of tamoxifen administration. (B) Fat and lean mass distribution in male WT and FAdora1^−/−^ mice before and after 10 weeks of high-fat diet feeding measured by EchoMRI. (C-J) Data were collected for male WT and FAdora1^−/−^ mice after 12-15 weeks of high-fat diet feeding. (C) Ambulatory activity for one full day/night cycle measured as area under the curve for total x-axis beam breaks in metabolic cages. (D) Respiratory exchange ratio (RER) in one full light and dark cycle. (E-G) Blood glucose (E), serum insulin (F), and serum NEFA (G) following an overnight fast (fasted) and 4-hour feeding period (refed). (H) Blood glucose during an i.p. insulin tolerance test with 0.75 IU/kg insulin. (I,J) Blood glucose and area under the curve (I), and serum insulin (J) during an i.p. glucose tolerance test with 1 g/kg glucose. Error bars represent SD. ****p<0.0001, t-test.

### A1R limits the lipolytic response to adrenergic stress in fasted but not refed mice

Adenosine levels are known to rise during stress, and adenosine signaling provides numerous protective benefits during stress responses (16,23,28). Stress-induced adrenergic stimulation activates lipolysis within adipose tissue, and exposure to high levels of NEFA during stress is associated with insulin resistance and lipotoxicity (18,19). We hypothesized that adenosine provides a protective limit on lipolysis in adipose tissue and that the loss of adipose A1R would result in a stronger lipolytic response to adrenergic stimulation and higher serum NEFA levels. To test this, we subjected WT and FAdora1^−/−^ mice to an isoproterenol tolerance test. After a short 6-hour fast to normalize serum NEFA levels, we administered 10 mg/kg isoproterenol by i.p. injection to stimulate adipocyte lipolysis. In WT mice, this dose of isoproterenol induces a modest and rapid increase in serum NEFA levels that returns to baseline levels within 2 hours (Fig. 4A). As we expected, treatment of FAdora1^−/−^ mice with isoproterenol resulted in a stronger stimulation of lipolysis achieving significantly higher NEFA levels, but also returning to baseline within 2 hours (Fig. 4A). The lipolytic response peaked 15 minutes after isoproterenol stimulation, which is not surprising considering the rapid induction of lipolysis and short half-life (~2 min) of isoproterenol (Fig. 4B).

**Figure 4.**
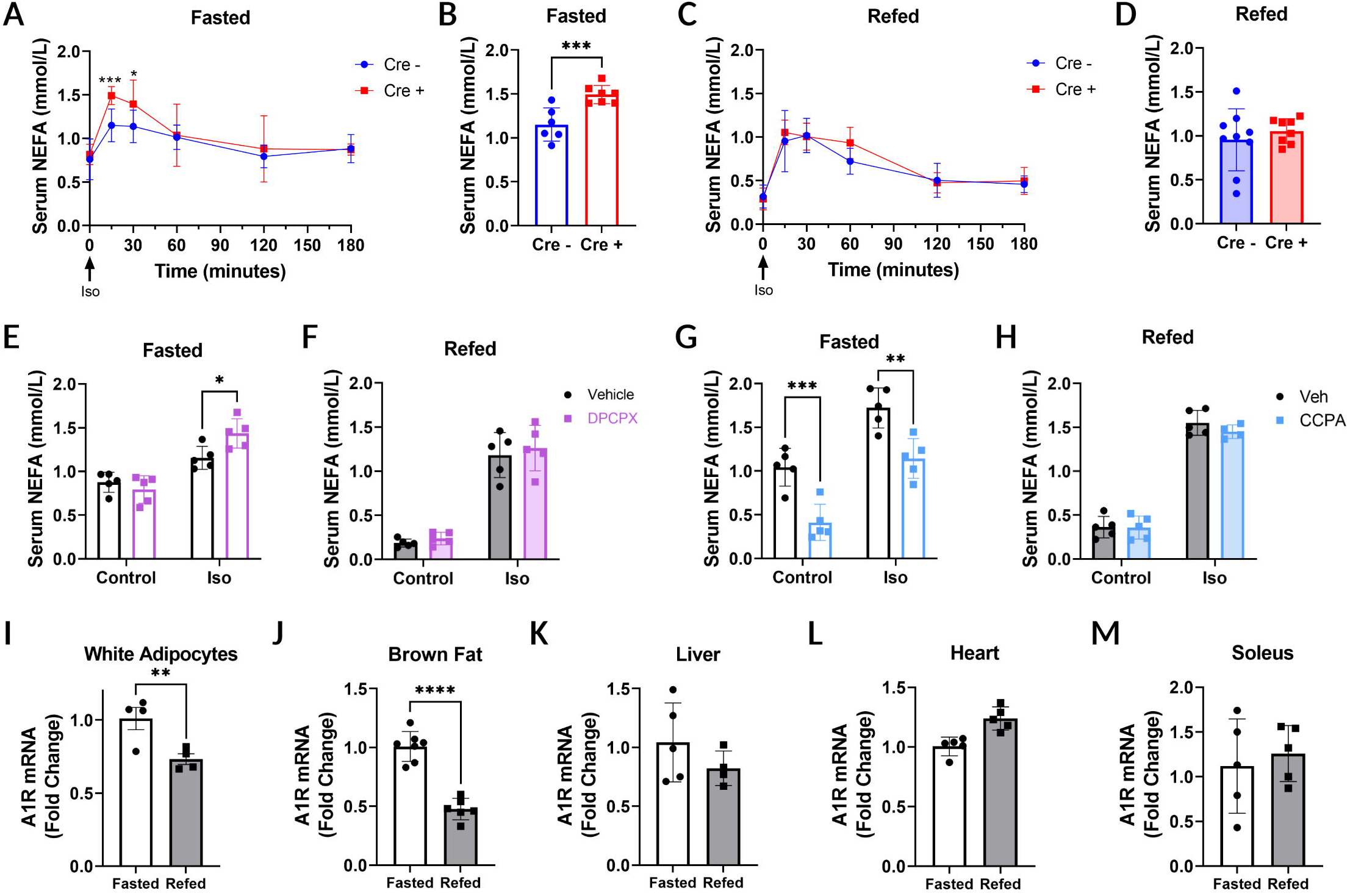
FAdora1^−/−^ mice are more lipolytically responsive in the fasted but not refed state due to A1R downregulation and desensitization after feeding. (A,B) Serum NEFA in 6-hour fasted male Cre negative WT (blue) and Cre positive FAdora1^−/−^ (red) mice at the indicated time points after i.p. injection of 10 mg/kg isoproterenol (A) or at 15 minutes after injection with isoproterenol (B). (C,D) Same as in (A,B) for mice that were fasted overnight and refed for 4 hours. (E) Serum NEFA in 6-hour fasted wild-type male C57BL/6J mice treated with vehicle (black) or 0.1 mg/kg DPCPX (purple) for 30 minutes both before (control) and 15 minutes after (iso) i.p. injection with 10 mg/kg isoproterenol. (F) Same as in (E) for mice that were fasted overnight and refed for 4 hours. (G,H) Same as in (E) and (F) except mice were treated with vehicle (black) or 0.5 mg/kg CCPA (light blue) for 30 minutes before isoproterenol injection. (I-M) Adipocytes and tissues from wild-type male C57BL/6J mice were harvested after either an overnight fast (white) or following a 4-hour refeeding period (shaded), and mRNA expression of A1R was measured in epididymal white adipocytes (I), brown adipose tissue (J), liver (K), heart (L), and soleus muscle (M). Error bars represent SD. *p<0.05, **p<0.01, ***p<0.001, ****p<0.0001, t-test or two-way ANOVA.

We also conducted these experiments under refed conditions by fasting mice overnight followed by a 4-hour refeeding period prior to stimulation with isoproterenol. As expected, both WT and FAdora1^−/−^ mice had significantly lower levels of serum NEFA following refeeding. However, there was no longer any difference in the elevation of NEFA following isoproterenol stimulation (Fig. 4C). Both WT and FAdora1^−/−^ mice reached a similar peak in serum NEFA at 15 minutes (Fig. 4D) and returned to baseline within 2 hours. Once again, no differences were seen in either the fasted or refed state with female FAdora1^−/−^ mice (Fig. S5A-D).

### Feeding desensitizes adipocytes to A1R signaling in vivo

The discrepancy between the fasted and refed state in FAdora1^−/−^ male mice suggested that refeeding may desensitize WT mice to signaling through A1R. To investigate this possibility in WT C57BL/6J mice, we used the same experimental setup as was used with the FAdora1^−/−^ mice in Fig. 4A–D; lipolysis was stimulated using isoproterenol in C57BL/6J mice that were either fasted for 6 hours or fasted overnight and refed for 4-hours. We first administered a 30-minute pre-treatment with the A1R antagonist 8-cyclopentyl-1,3-dipropylxanthine (DPCPX) and collected serum to assess NEFA levels under control conditions before subsequent induction of lipolysis with isoproterenol for 15 minutes. Treatment with DPCPX alone did not alter NEFA levels, however DPCPX-treated mice exhibited a stronger lipolytic response to isoproterenol compared with vehicle-treated mice under fasted conditions (Fig. 4E). This difference in lipolytic response was not present in refed mice, consistent with the results seen in FAdora1^−/−^ mice (Fig. 4F).

While these data alongside the results obtained with FAdora1^−/−^ mice suggested a loss of sensitivity to A1R signaling in response to feeding, it is possible the discrepancy in these phenotypes between the fasted and refed conditions could be explained by differences in endogenous adenosine levels. A substantial reduction in extracellular adenosine following refeeding would also produce a lack of any phenotype with both genetic knockout and antagonism of A1R. Therefore, we subjected fasted and refed C57BL/6J mice to pre-treatment with the A1R agonist 2-chloro-N^6^-cyclopentyladenosine (CCPA) and isoproterenol stimulation. Under fasted conditions, agonism of A1R with CCPA produced a significant decrease in serum NEFA in both the control and isoproterenol stimulated states (Fig. 4G). Strikingly, all effects of A1R agonism on serum NEFA were eliminated under refed conditions, even with isoproterenol stimulation (Fig. 4H). These data confirmed that adipocytes are indeed desensitized to A1R signaling in response to feeding, and that changes in extracellular adenosine are unlikely to be responsible for reduced signaling through A1R during the fed state.

### A1R is acutely downregulated in response to feeding specifically in adipocytes

To better understand the mechanism driving A1R desensitization, we subjected wild-type C57BL/6J mice to either an overnight fast alone or an overnight fast followed by a 4-hour refeeding period and measured A1R mRNA expression levels. We found that in isolated epididymal white adipocytes there was a significant reduction in A1R mRNA levels in refed mice (Fig. 4I). We saw a similar suppression of A1R expression in brown adipose tissue (Fig. 4J), but not in liver, heart, or skeletal muscle (Fig. 4K–M). Thus, feeding induces a rapid downregulation of A1R mRNA expression specifically within adipocytes.

### Insulin regulates A1R transcription in adipocytes via the PI3K-Akt pathway

To further investigate the signaling mechanisms which regulate A1R transcription in response to feeding, we used 3T3-L1 adipocytes with an overnight serum starve to approximate adipocytes under fasted conditions. A 4-hour treatment with 10% fetal bovine serum (FBS) to simulate feeding had little effect on A1R transcription (Fig. 5A). However, in adipocytes insulin is perhaps the most important anabolic hormone and the amount of insulin present in media containing 10% FBS is less than 10 pmol/L according to the certificate of analysis provided for the lot number used. The introduction of additional insulin into the media in the presence or absence of serum produced a drastic decrease in A1R mRNA (Fig. 5A). This effect was dose-dependent, with an EC_50_ well within the physiological range at ~0.15 nmol/L (~25 μU/mL). This effect was also rapid, with near maximal reduction in mRNA within 2 hours and sustained for at least 8 hours in the presence of insulin (Fig. 5B).

**Figure 5.**
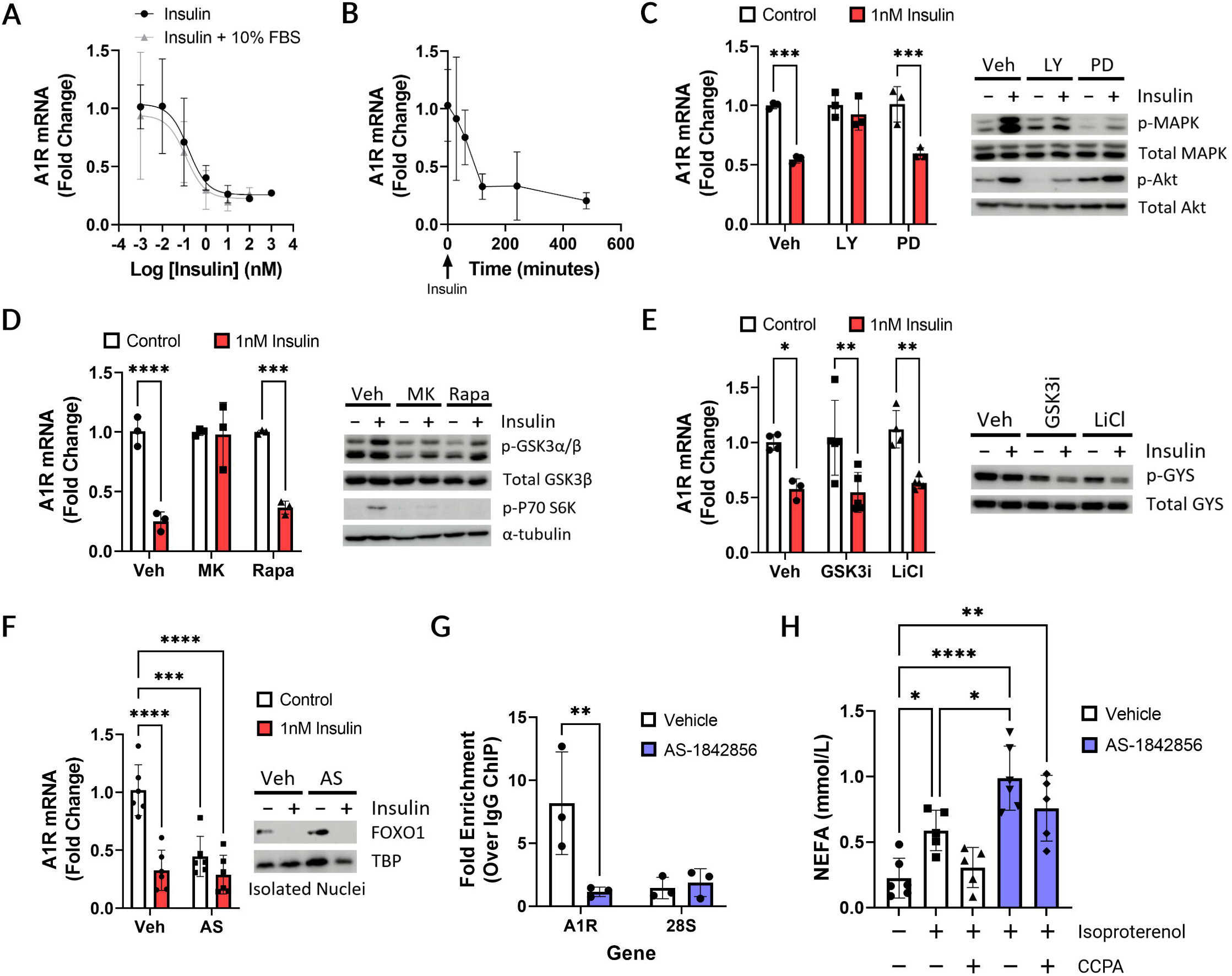
The insulin-Akt-FOXO1 pathway acutely regulates A1R expression to modulate adenosine signaling in adipocytes. (A-F) 3T3-L1 adipocytes 10-12 days after differentiation were serum starved overnight and treated with insulin as indicated for either 4 hours (mRNA analysis) or 15 minutes (phospho-protein western blot analysis). (A) Dose response of insulin effects on A1R expression in the presence or absence of 10% FBS. (B) Time course for the suppression of A1R expression by 10 nmol/L insulin. (C) Effect of vehicle (white) or 1 nmol/L insulin (red) on A1R mRNA expression, phospho-MAPK (T202/Y204), and phospho-Akt (S473) after 15-minute pre-treatment with vehicle (Veh), 50 μmol/L LY294002 (LY), or 25 μmol/L PD98059 (PD). (D) Effect of 1 nmol/L insulin on A1R mRNA expression, phospho-GSK3α/β (S21/S9), and phospho-P70 S6K (T389) after 15-minute pre-treatment with vehicle (Veh), 25 μmol/L MK-2206 (MK), or 250 nmol/L rapamycin (Rapa). (E) Effect of 1 nmol/L insulin on A1R mRNA expression and phospho-GYS (S641) after 15-minute pre-treatment with vehicle (Veh), 1.4 μmol/L GSK3 inhibitor IX (GSK3i), or 30 mmol/L LiCl. (F) Effect of 1 nmol/L insulin on A1R mRNA expression and nuclear localization of FOXO1 after 15-minute pre-treatment with vehicle (Veh) or 1 μmol/L AS-1842856 (AS). (G) Fold-enrichment by ChIP with FOXO1 over normal rabbit IgG of A1R promoter region (A1R) or 28S rDNA control region (28S) after 4-hour treatment of 3T3-L1 adipocytes with vehicle (white) or 1 μmol/L AS1842856 (blue). (H) NEFA released into the media by isolated adipocytes that were treated with vehicle or 1 μmol/L AS-1842856 for 4 hours, followed by vehicle or 100 nmol/L CCPA for 15 minutes in the presence of adenosine deaminase and a subsequent 30-minute treatment with vehicle or 30 nmol/L isoproterenol. Error bars represent SD. *p<0.05, **p<0.01, ***p<0.001, ****p<0.0001, one-way or two-way ANOVA.

We next sought to determine the signaling mechanisms downstream of insulin that are responsible for A1R downregulation. The insulin receptor signaling pathway has numerous effectors but can be simplified to two immediate primary effectors: phosphoinositide 3-kinase (PI3K) and mitogen activated protein kinase kinase (MAP2K). To interrogate these pathways, we utilized the pharmacological inhibitors LY294002 and PD98059 to inhibit PI3K and MAP2K respectively. While a 4-hour treatment with insulin alone suppressed A1R transcription and strongly activated the PI3K pathway as seen with phosphorylation of Akt at S473, a 30-minute pretreatment with LY294002 prevented both effects (Fig. 5C). Meanwhile, inhibition of MAP2K with PD98059 significantly blunted insulin-induced phosphorylation of P42/44 MAPK at T202/Y204, but did not prevent the suppression of A1R transcription.

Downstream of PI3K, the protein kinase Akt is an important intermediary effector that phosphorylates proteins important for insulin signaling in adipocytes, such as mTORC1 and GSK3β. MK-2206 is a specific inhibitor of Akt1/2/3 which was able to prevent both insulin-induced phosphorylation of GSK3α/β at S21/9 and suppression of A1R (Fig. 5D). Treatment with rapamycin, a specific inhibitor of mTORC1, blocked phosphorylation of the downstream target P70 S6 kinase (S6K) at T389, however there was no impact to the suppression of A1R by insulin (Fig. 5D). We also inhibited the action of GSK3 with the GSK3 inhibitor IX, and with LiCl. Both treatments significantly impaired the phosphorylation of glycogen synthase (GYS) at S641, but neither had any impact on the suppression of A1R (Fig. 5E). Thus, A1R transcription is suppressed by insulin via the PI3K-Akt pathway but is not dependent on mTORC1 or GSK3β.

### A1R transcription is regulated by FOXO1

The forkhead box proteins within the FOXO family are transcription factors with known sensitivity to nutrient status. FOXO1 is highly expressed in insulin sensitive tissues such as adipose and liver, and contains 3 known phosphorylation sites that are targets of Akt (29). Phosphorylation of FOXO1 by Akt inhibits its transcriptional activity by promoting the interaction of FOXO1 with 14-3-3 proteins and retention within the cytoplasm (30). Further, both FOXO1 and A1R are involved in adipocyte differentiation and adipogenesis and show a similar pattern of upregulation as adipocyte differentiation progresses (31,32). We therefore hypothesized that FOXO1 may be directly regulating A1R transcription in adipocytes. To explore this hypothesis, we utilized the pharmacological inhibitor AS-1842856 which competitively inhibits the DNA binding of FOXO1 (33). Treatment of 3T3-L1 adipocytes with insulin both suppressed A1R transcription and induced the translocation of FOXO1 out of the nucleus as determined by western blot of isolated nuclei (Fig. 5F). Treatment with AS-1842856 alone resulted in the suppression of A1R transcription to a similar level as insulin treatment but did not affect FOXO1 localization. Finally, treatment with both insulin and AS-1842856 did not result in any further suppression of A1R, suggesting that insulin and AS-1842856 suppress A1R transcription through a similar mechanism.

We next wanted to confirm that FOXO1 directly binds to the promoter of A1R to regulate its transcription. We used the LASAGNA-search 2.0 tool to identify sites containing the consensus motif for FOXO1 binding within the A1R promoter (34). We identified a region within the first intron of the A1R gene locus containing 4 consensus motifs within a span of 50 bp. We next used chromatin immunoprecipitation (ChIP) to enrich for FOXO1 binding sites, followed by qPCR and found this region within the A1R locus was significantly enriched compared with a normal IgG control ChIP (Fig. 5G). There was no enrichment in the control region within 28S rDNA.

### Inhibition of FOXO1 in adipocytes induces A1R insensitivity

Thus far, we have shown that feeding suppresses A1R transcription in adipocytes, and that insulin is capable of suppressing A1R transcription through the phosphorylation and inactivation of FOXO1. We therefore wondered what the effects of direct FOXO1 inhibition would be on adipocyte lipolysis and A1R sensitivity. We isolated adipocytes from the epididymal fat pads of overnight fasted male C57BL/6J mice and treated them with either vehicle or the FOXO1 inhibitor AS-1842856 for 4 hours. We then treated the cells with vehicle or the A1R agonist CCPA in the presence of adenosine deaminase for 15 minutes followed by a 30-minute treatment with vehicle or isoproterenol and measured NEFA release into the media. In the absence of FOXO1 inhibition, isoproterenol strongly increased the concentration of NEFA within the media and the addition of CCPA suppressed NEFA back to control levels as expected (Fig. 5H). While the inhibition of FOXO1 alone had no effect on NEFA release (not shown), the addition of isoproterenol in these adipocytes resulted in a significantly higher induction in lipolysis which CCPA was not able to suppress (Fig. 5H). Taken together, these data demonstrate that A1R transcription is acutely regulated via the insulin-Akt-FOXO1 signaling axis within adipocytes, and that insulin stimulation results in a rapid downregulation of A1R transcription and reduced sensitivity to A1R signaling.

### Obesity disrupts A1R regulation of lipolysis in both the fasted and refed state

Changes in adenosine signaling have been previously implicated in the development of insulin resistance in obesity. Multiple groups have demonstrated that obesity correlates with increased adenosine levels in human adipose tissue, yet reduced expression of A1R in adipose tissue in humans (35,36). We wanted to know how obesity impacts not only A1R signaling in adipocytes, but also the regulation of A1R expression by feeding. We first placed C57BL/6J mice on either a normal chow or high fat diet (HFD) for 12 weeks, after which normal chow mice weighed on average 29.4 g ± 1.7 g and HFD mice weighed 46.5 g ± 3.2 g. We then compared A1R mRNA expression in isolated adipocytes from overnight fasted mice and confirmed that HFD also results in significantly reduced expression of A1R mRNA in mice (Fig. 6A). When A1R expression between fasted and refed HFD mice is compared, we found that feeding no longer suppressed A1R transcription (Fig. 6B).

**Figure 6.**
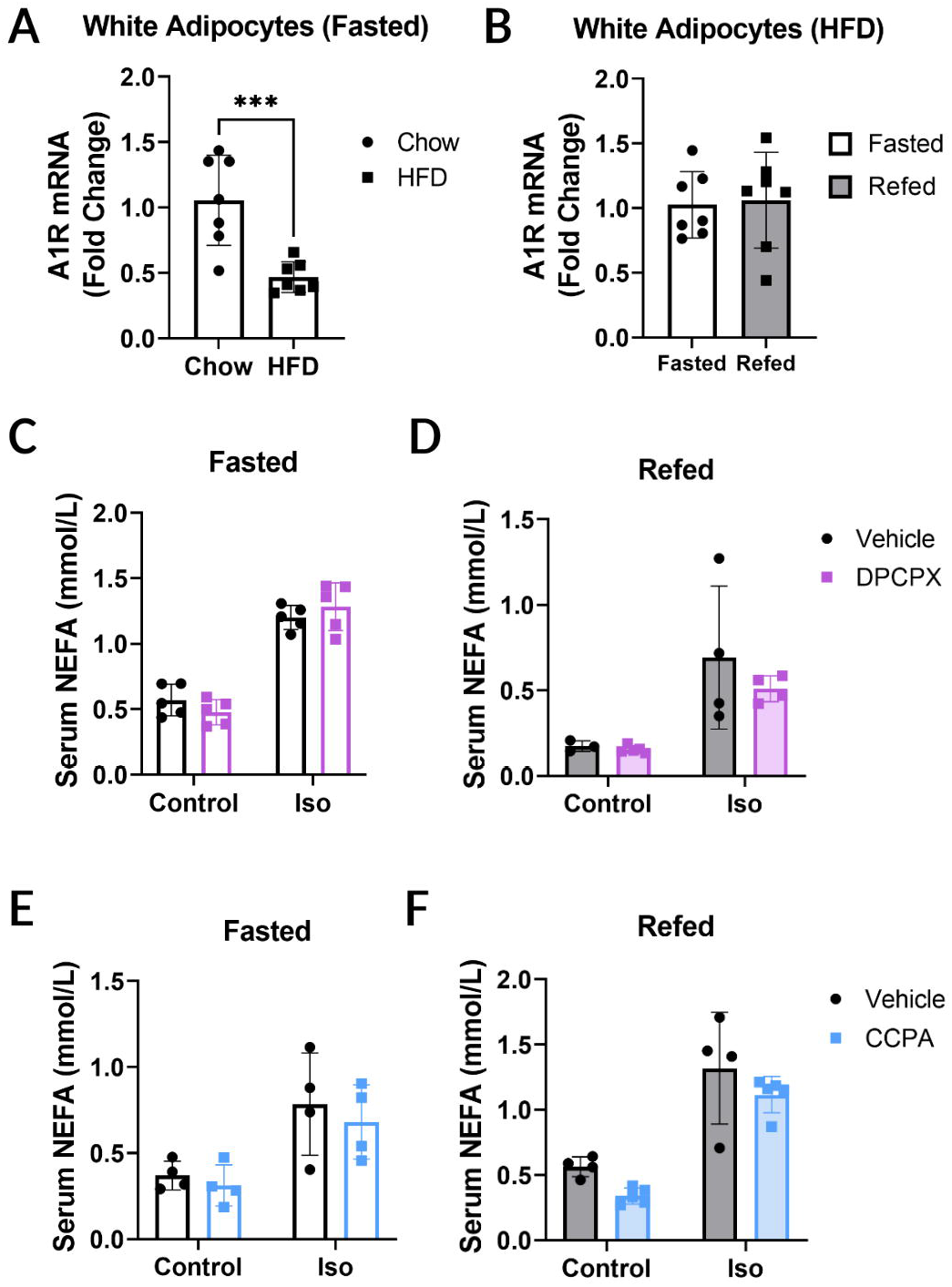
High-fat diet downregulates and desensitizes A1R in mouse adipocytes and disrupts regulation of A1R expression by feeding. (A) mRNA expression of A1R in epididymal white adipocytes from male wild-type C57BL/6J mice fed either normal chow control diet (Chow) or high-fat diet (HFD) for 12 weeks following an overnight fast. (B) mRNA expression of A1R in epididymal white adipocytes from male wild-type C57BL/6J mice fed high-fat diet for 12 weeks following either an overnight fast or a 4-hour refeeding period. (C-F) Male wild-type C57BL/6J mice were fed high-fat diet for 12 weeks before experimentation. (C) Serum NEFA in 6-hour fasted mice treated with 0.1 mg/kg DPCPX for 30 minutes both before (control) and 15 minutes after (iso) i.p. injection with 10 mg/kg isoproterenol. (D) Same as in (C) for mice that were fasted overnight and refed for 4 hours. (E,F) Same as in (C) and (D) except mice were treated with 0.5 mg/kg CCPA for 30 minutes before isoproterenol injection. Error bars represent SD. ***p<0.001, t-test.

We next repeated the experiments using the A1R antagonist DPCPX alongside isoproterenol stimulation of lipolysis with mice on HFD. We found that DPCPX did not result in an increased lipolytic response to isoproterenol under either fasted or refed conditions in these mice (Fig. 6C and D). Further, administration of the A1R agonist CCPA was also unable to suppress lipolysis under either fasted or refed conditions in HFD mice (Fig. 6E and F). These results demonstrate that, in addition to suppressing A1R transcription, obesity disrupts the regulation of A1R transcription by fasting and feeding and the regulation of lipolysis by A1R signaling.

## Discussion

In this study, we investigated the role of the A_1_ adenosine receptor in adipose tissue in the regulation of lipid and glucose metabolism. Our findings agree with previous studies in isolated adipocytes and show that A1R augments insulin suppression of lipolysis and limits the lipolytic response to adrenergic stress *in vivo*. Further, this study agrees with findings from mice with A1R overexpression using the aP2 promoter (37), and demonstrates that A1R action in adipose tissue serves to protect mice from obesity-induced glucose intolerance. Moreover, we found that loss of adipose A1R only resulted in a higher lipolytic response to stress under fasted conditions. This led to the key finding of this study that elevated insulin levels in the fed state rapidly downregulate and desensitize A1R specifically in adipocytes, and this reduction is mediated by the transcription factor FOXO1.

The regulation of A1R by FOXO1 initially presents as counterintuitive since FOXO1 drives high expression of A1R in the fasted state when conditions are primarily catabolic, while signaling through A1R is primarily anabolic. However, this view of A1R in steady state misses a critical component of this regulation, which is the lag between stimulation by insulin and an observed decrease in A1R mRNA. Considering the data presented in this study, we propose two distinct roles for A1R signaling in adipocytes which are not mutually exclusive. First, we propose that active FOXO1 drives high A1R in the fasted state to limit deleterious effects such as lipotoxicity resulting from the overstimulation of lipolysis by catabolic signals (Fig. 7, top left). Second, we propose that during the anabolic state A1R promotes insulin action to suppress lipolysis (Fig. 7, top right), but a postprandial decline in A1R levels mediated through reduced FOXO1 transcriptional activity (Fig. 7, bottom right) facilitates the reentrance of adipocytes into the catabolic state by removing an inhibitor of lipolysis (Fig. 7, bottom left). Homeostasis posits that an organism responds to changing conditions to maintain the internal milieu constant, while allostasis suggests that organisms alter the internal milieu in order to meet anticipated demands through neuronal mechanisms (46,47). It has been suggested that adenosine plays a more significant role in allostasis than in homeostasis, and the decline in A1R expression appears to be an allostatic adaptation to the predicted eventual shift from anabolism to catabolism (45).

**Figure 7.**
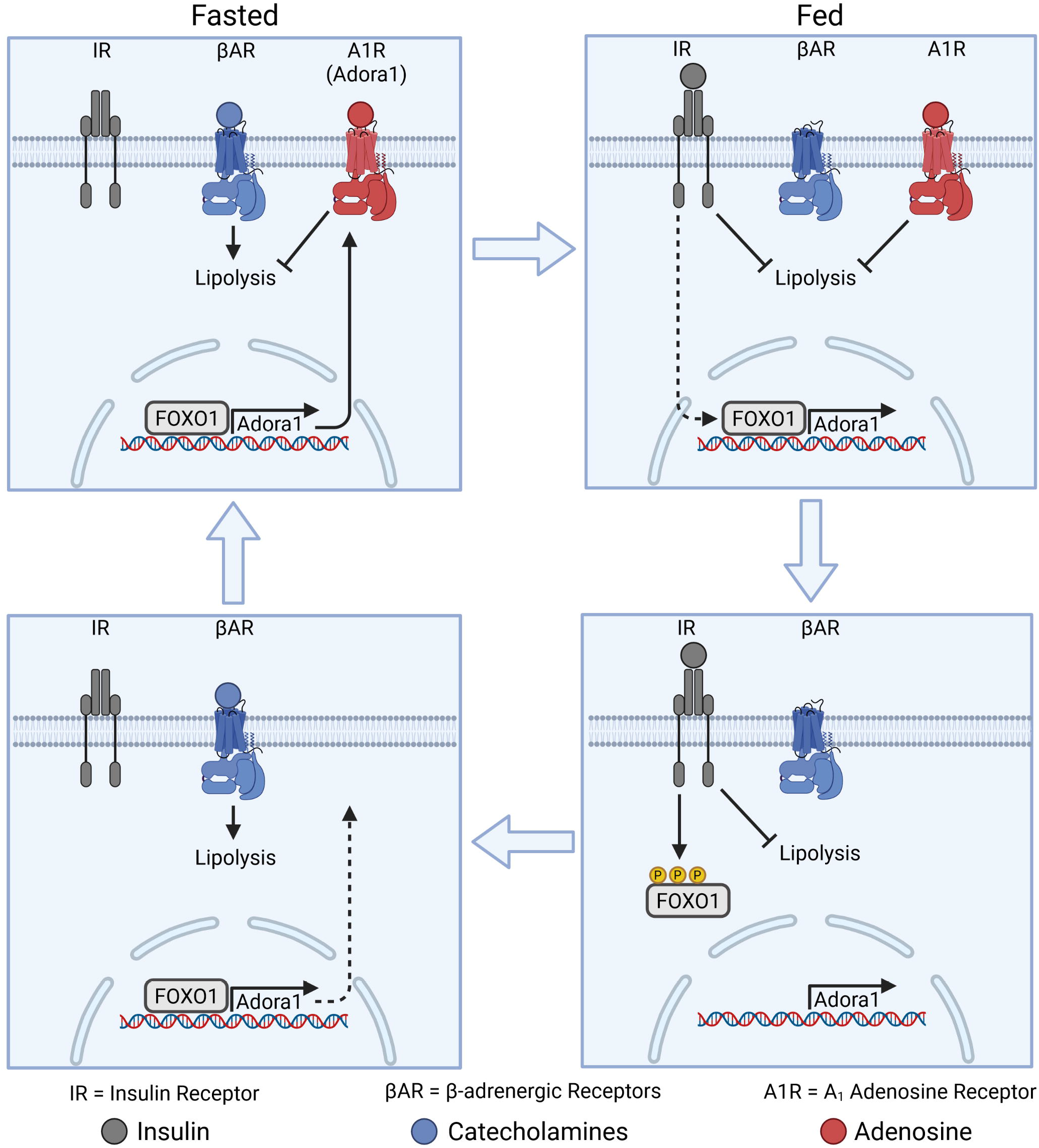
Proposed mechanism for FOXO1 modulation of adenosine signaling in adipocytes during fasting and feeding. Under fasted conditions, active FOXO1 drives high expression of A1R which serves to limit lipolysis stimulated by catecholamines during stress. Upon feeding, catecholamine levels decrease while insulin levels increase. Insulin acts to suppress lipolysis and induces the phosphorylation and inactivation of FOXO1, decreasing transcription of A1R. Initially, A1R expression persists at high levels and signaling through A1R augments the suppression of lipolysis by insulin. High insulin levels and inactive FOXO1 under fed conditions eventually lead to reduced A1R expression and the removal of the inhibitory effect of A1R on lipolysis. Reduced nutrient levels brought on by fasting lead to decreased insulin and increased catecholamines. Lower expression of A1R facilitates the initial induction of lipolysis by catecholamines upon fasting. Decreased insulin levels lead to active FOXO1 which begins to increase A1R to reengage its limitation on lipolysis in the fasted state.

The role of A1R in adipocytes *in vitro* has been well characterized by many groups who have demonstrated that adenosine enhances insulin-stimulated glucose uptake and lipogenesis and suppresses basal and stimulated lipolysis within adipocytes (12,15,38–41). Investigating the role of adipose A1R *in vivo*, however, has been challenging as A1R is also involved in the regulation of numerous other tissues such as the brain, heart, skeletal muscle, and pancreatic islet cells. This has led to conflicting results with glucose metabolism in whole-body knockouts of A1R, where they have been shown to display impaired glucose tolerance (9), improved glucose tolerance (7), and no change to glucose tolerance (10). Because all three groups saw changes in body weight and the secretion of glucagon and insulin, the role of adipose tissue A1R in these contexts has remained unclear. Dong and colleagues provided good evidence that adipose A1R improves glucose tolerance and insulin sensitivity by overexpressing A1R using the aP2 promoter (37). These mice did not display the previously seen changes in body weight or insulin levels, however overexpression clearly enhanced the tonic activity of A1R resulting in suppression of basal lipolysis and reduced serum NEFA levels. We now show that endogenous A1R in adipose also promotes glucose tolerance and insulin sensitivity. Our studies show that mice with an inducible, adipocyte-specific knockout of A1R are similarly devoid of the changes in body weight and insulin and glucagon secretion, while also lacking any changes to basal fasted and fed serum NEFA levels. Instead, we saw that loss of adipose A1R impairs the kinetics of insulin-mediated suppression of lipolysis rather than the final homeostatic set points. FAdora1^−/−^ mice also developed glucose intolerance when placed on HFD, without changes to insulin secretion or sensitivity. The suppression of hepatic glucose production by insulin has been shown to be dependent on the appropriate suppression of adipocyte lipolysis (26,27). Therefore, it is likely this glucose intolerance is driven by increased endogenous glucose production within FAdora1^−/−^ mice, consistent with their impaired ability to reduce serum NEFA in response to insulin.

Adenosine levels are increased in response to adrenergic stress, and adenosine action on A1R suppresses adrenergic stimulation of lipolysis in adipocytes (13,16,41). A1R agonists have been clearly demonstrated to suppress basal and ß-adrenergic-stimulated lipolysis *in vivo*, however these experiments were performed in fasted subjects, even in human patients, as is typical for most metabolic studies (25,42–44). While we found that both genetic loss of adipose A1R and general antagonism of A1R resulted in a stronger lipolytic response to adrenergic stimuli in fasted conditions, this effect was not present in refed mice. We did not measure the effects of feeding on interstitial or local plasma adenosine levels, however the desensitization of adipose A1R under refed conditions was independent of any potential reduction in adenosine as lipolysis in WT refed mice was also unaffected by a high dose (0.5 mg/kg) of the full A1R agonist CCPA. It is also unlikely that alternative pathways influencing cAMP levels, such as insulin activation of phosphodiesterases, are confounding our results, as isoproterenol still produced a clear induction in lipolysis within refed mice which was not suppressed by A1R agonism. We did not, however, probe the effects of feeding on Gαi proteins, and it may be interesting to investigate the sensitivity of adipocytes to other antilipolytic agents, such as nicotinic acid or prostaglandin E_2_, under refed conditions.

We found that the loss in adipose A1R sensitivity following refeeding can at least partially be explained by acute downregulation of A1R mRNA. We would like to note, however, that the reductions in mRNA measured *in vivo* may not be large enough to account for the full loss of A1R sensitivity observed, and other mechanisms such as post-translational modifications or changes in Gαi proteins may be involved. Still, a significant reduction in A1R mRNA was mediated by the insulin-Akt-FOXO1 signaling axis, and this relationship between FOXO1 and A1R may explain a central shift in adenosine receptors that occurs during adipogenesis. Gharibi et. al. demonstrated that A1R promotes adipogenesis, and that A1R expression is dramatically increased several days into the adipocyte differentiation process before falling to a level that is still significantly elevated over preadipocytes (32). Interestingly, Nakae and colleagues demonstrated that FOXO1 also promotes adipogenesis, and FOXO1 expression follows a strikingly similar pattern and timing during adipocyte differentiation (31). It follows that increased expression of A1R during the differentiation process is driven by FOXO1, although further work is needed to demonstrate this mechanism.

Obesity has been shown to reduce responsiveness to inhibitors of cyclic AMP accumulation such as adenosine (36). This has been associated with reduced expression of A1R in adipocytes isolated from obese human patients as compared to lean patients (35,36). We observed that A1R expression is also reduced in adipocytes isolated from obese mice and confirmed that this leads to a lack of A1R sensitivity in obese mice under both fasted and refed conditions. It is possible that both reduced A1R expression and the loss of cyclical fluctuations in expression between fasting and feeding may contribute to insulin resistance and glucose intolerance.

In summary, adipose A1R is acutely regulated by feeding to modulate the adenosine response and allow adipose tissue to adequately adapt to changing nutrient conditions. High expression of adipose A1R under fasted conditions is necessary to limit adrenergic-driven lipolysis and enhance insulin-mediated suppression of lipolysis.

## Acknowledgments

The authors would like to thank Dr. Stanislav Zakharenko for sharing the floxed *Adora1* mice with us. We also thank Dr. Ira G. Schulman and Dr. Mohan Manjegowda for their input and resources in chromatin immunoprecipitation experiments. This data has been previously presented at the Mid-Atlantic Diabetes and Obesity Research Symposium and published on BioRxiv. Figure 7 was made with Biorender.

## Author Contributions

M.E.G. and S.R.H. assessed knockout and performed metabolic and lipolytic characterization of mice and performed qPCR on mouse tissues. S.R.H. performed metabolic caging studies. M.E.G. and M.A.L. performed capadenoson experiments. M.E.G. and M.C.L. performed 3T3-L1 studies. M.E.G., S.R.H. and T.E.H. performed isolated adipocyte studies. I.M.B. provided significant conceptual and technical input on ChIP experiments. T.E.H., J.L., and M.E.G. conceived the study and designed the experiments. M.E.G. and T.E.H. wrote the manuscript. M.E.G. is the guarantor of the work and, as such, had full access to all the data in the study and takes responsibility for the integrity of the data and the accuracy of the data analysis.

