## Supplemental Material for "Feeding desensitizes A1 adenosine receptors in adipose through FOXO1-mediated transcriptional regulation"

### Materials and Methods

#### *Materials*

All pharmacological compounds were obtained from Cayman Chemical (Ann Arbor, MI) unless otherwise stated. Capadenoson was obtained from MedChemExpress (Monmouth Junction, NJ). 3-isobutyl-1-methylxanthine (IBMX), dexamethasone, GSK3 inhibitor IX, formaldehyde, and peanut oil were obtained from Sigma-Aldrich (St. Louis, MO). Regular human insulin (Humulin R) was obtained from Eli Lilly (Indianapolis, IN). All antibodies were obtained from Cell Signaling (Danvers, MA) unless otherwise specified. Adora1 antibody (PA1-041a) was obtained from ThermoFisher (Waltham, MA). The FOXO1 antibody (18592-1-AP) used for ChIP-qPCR was obtained from ProteinTech (Rosemont, IL).

#### *Animals*

AdipoQ-cre-ERT2 (C57BL/6-Tg(Adipoq-cre/ERT2)1Soff/J, Jax #025124) and C57BL/6J (Jax #000664) mice were from the Jackson Laboratory. In FAdora1$^{-/-}$ mice, knockout was induced at 8-weeks by daily injection of tamoxifen (50 mg/kg) dissolved in peanut oil for 10 days, followed by a 14-day washout period prior to experimentation. Cre-negative wild-type control mice were age and sex-matched littermates that received the same tamoxifen treatment protocol. For high-fat diet (HFD) studies in WT and FAdora1$^{-/-}$ mice, animals were placed on diet for 12 weeks starting at 8 weeks of age (BioServ, F1850). For diet studies in wild-type animals, C57BL/6J mice at Jackson Labs were put on HFD beginning at 6 weeks of age (#380050) and were shipped to the University of Virginia vivarium at 14 weeks of age. The mice were then maintained on HFD

(Research Diets, D12492) for another four weeks before experiments, for a total of 12 weeks of HFD.

*Animal Procedures*

To measure insulin and NEFA levels, blood was collected from the tail vein of mice, and serum was prepared and frozen at -80°C until analysis. Glucose was measured in tail vein blood using a One-touch Ultra glucometer. Heart rate was measured by electrocardiogram in mice anesthetized with isoflurane. Insulin, glucose, and all pharmacological compounds were administered via i.p. injection in 0.9% NaCl, except for glucose which was administered as a 20% glucose solution. Body weight was measured once weekly at the same time of day for both normal chow and HFD mice. Ambulatory activity, carbon dioxide production ($VCO_2$), and oxygen consumption ($VO_2$) were measured using an Oxymax metabolic chamber system (Comprehensive Laboratory Animal Monitoring System) from Columbus Instruments (Columbus, OH). One full day and night was allowed for acclimation. Mice were either fasted overnight (fasted) or then given access to chow and softened food for 4 hours (refed) as indicated, with exception of the fasted condition for the isoproterenol tolerance tests where mice were fasted for 6 hours. For tissue harvesting, mice were euthanized by cervical dislocation, and tissues were flash frozen in liquid nitrogen. For primary adipocyte isolations, epididymal adipose tissue was not frozen and was processed immediately.

To measure the effects of capadenoson on lipolysis and heart rate, mice were fasted overnight and administered with vehicle or 1 mg/kg capadenoson i.p. Serum was collected and heart rate measured 2 hours after administration.

Glucose tolerance tests were performed following an overnight fast. Mice were injected i.p. with 20% glucose solution (1 g/kg), and glucose was measured and serum collected at the indicated time points. Insulin tolerance tests were performed in the morning in *ad libitum* fed mice. Mice were injected i.p. with insulin (0.75 IU/kg) in 0.9% NaCl and glucose was measured at the indicated time points.

For fasted and refed measurements, mice were fasted overnight, glucose was measured and serum was collected and frozen for later analysis of insulin and NEFA. Mice were then given *ad libitum* access to food softened with water for 4 hours and measurements were repeated.

To measure the suppression of lipolysis by insulin, mice were fasted overnight and administered insulin i.p. (0.75 IU/kg). Serum was collected at 0 and 15 minutes after insulin administration. Mice were monitored for an additional 1 hour for hypoglycemia and administered glucose as needed.

To measure lipolytic responses in an isoproterenol tolerance test, WT and FAdora1$^{-/-}$ mice were administered 10 mg/kg isoproterenol via i.p. injection, and serum was collected at the indicated time points. C57BL/6J mice were administered vehicle, 0.1 mg/kg DPCPX, or 0.5 mg/kg CCPA via i.p. injection 30 minutes prior to isoproterenol administration. Serum was collected at 0 (control) and 15 (iso) minutes after isoproterenol administration.

*Adipocyte isolation*

Adipocytes were isolated as previously described (1,2). Epididymal fat pads were minced in low phosphate buffer (145 mmol/L NaCl, 5.4 mmol/L KCl, 1.4 mmol/L $CaCl_2$, 1.4 mmol/L $MgSO_4$, 0.2 mmol/L $NaH_2PO_4$, 5 mmol/L glucose, 10 mmol/L HEPES, pH 7.4) containing 2.5% bovine serum albumin (BSA, Sigma-Aldrich, St. Louis, MO), 3 mg/mL collagenase type I (Worthington, Lakewood, NJ), and 100 nmol/L adenosine, and digested for 45 minutes in a 37°C water bath with mild agitation. For protein and RNA extraction, adipocytes were washed twice with buffer containing 0.1% BSA, and once with buffer without BSA. For lipolysis assays, adipocytes were washed three times with buffer containing 2.5% BSA.

*qPCR Primers*

Primers for *Adora1*, *Adora2A*, *Adora2B*, and *Adora3* were sourced from the PrimerBank of the Center for Computational and Integrative Biology of Massachusetts General Hospital and Harvard University, and were used to determine changes in mRNA expression for all experiments with wildtype mice and 3T3-L1 adipocytes (3–5). Primers targeting the 3' terminal exon for *Adora1* (exon 3) were designed using Primer3, and were used to determine the efficiency of the tamoxifen-induced knockout in FAdora1$^{-/-}$ mice (6,7).

| Target | Forward Primer | Reverse Primer |
|---|---|---|
| *Ppia* | CGATGACGAGCCCTTGG | TCTGCTGTCTTTGGAACTTTGTC |
| *Adora1* | TGTGCCCGGAAATGTACTGG | TCTGTGGCCCAATGTTGATAAG |
| *Adora2A* | GCCATCCCATTCGCCATCA | GCAATAGCCAAGAGGCTGAAGA |
| *Adora2B* | AGCTAGAGACGCAAGACGC | GTGGGGGTCTGTAATGCACT |
| *Adora3* | AAGGTGAAATCAGGTGTTGAGC | AGGCAATAATGTTGCACGAGT |
| *Adora1* (exon 3) | ACTTCTTCGTCTGGGTGCTG | TCCCGTAGTACTTCTGGGGG |

*Culture and treatment of 3T3-L1 adipocytes*

3T3-L1 fibroblasts and adipocytes (ATCC) were cultured and differentiated as previously reported (1). Briefly, 3T3-L1 fibroblasts were maintained and grown to confluency in DMEM (Gibco, 11965-092) with 10% newborn calf serum (Gibco, 16010-159), 1% fetal bovine serum (FBS, Gemini Bio Products, Lot #A15H74K), and 1% antibiotic-antimycotic (AA, Gibco). Cells were differentiated 3 days after becoming confluent in DMEM with 10% FBS, 1% AA, and containing 0.25 IU/mL insulin, 0.5 mmol/L IBMX, 0.25 μmol/L dexamethasone, and 10 μmol/L pioglitazone. After 4 days, adipocytes were maintained in DMEM with 10% FBS.

Experiments were performed with 3T3-L1 adipocytes 10 – 12 days after differentiation. Adipocytes were serum starved overnight in DMEM with 0.25% FBS and 0.25% BSA and treated with the indicated inhibitors at the concentrations described in figure legends 30 minutes prior to the addition of insulin. For qPCR analysis and FOXO1 translocation western blots, cells were treated with 1 nmol/L insulin for 4 hours unless otherwise indicated. For western blot analysis of phospho-proteins, cells were treated with 1 nmol/L insulin for 15 minutes.

*Sample preparation and antibodies for western blots*

Adipocytes were lysed with cell lysis buffer (10 mmol/L $Na_2HPO_4$, 50 mmol/L β-glycerophosphate disodium salt hydrate, 50 mmol/L NaF, 1 mmol/L EDTA, 1 mmol/L EGTA, pH 7.4) supplemented with protease inhibitors and 0.5 mmol/L dithiothreitol, homogenized by shearing through a 22G needle, and cleared by centrifugation at 17,000 x *g* for 10 minutes at 4°C. Tissues were homogenized by Potter-Elvehjem tissue grinder in Tissue Protein Extraction Reagent (Thermo) supplemented with protease inhibitors and 0.5 mmol/L dithiothreitol and cleared by centrifugation at 17,000 x *g* for 10 minutes at 4°C. Protein concentrations were determined using BCA assay (Pierce). Western blots were probed as indicated with β-actin (#A228, Sigma), α-tubulin (T9026, Sigma), GAPDH (#5174), TBP (#8515), A1R (#PA1-041a, ThermoFisher), phospho-Akt (S473, #9271), phospho-MAPK (T202/Y204, #9101), phospho-GSK3α/β (S21/S9, #9331), phospho-P70 S6K (T389, #9205), phospho-GYS (S641, #3891), total Akt (#2920), total P42/P44 MAPK (#4696), total GSK3β (#9315), total GYS (#3893), and FOXO1 (#2880).

*Nuclear isolation*

3T3-L1 adipocytes were washed twice with cold PBS and lysed in sucrose buffer (250 mmol/L sucrose, 50 mmol/L NaF, 1 mmol/L EDTA, 50 mmol/L Tris, pH 7.4) with Dounce homogenization. Lysates were incubated on ice for 5 minutes, and then centrifuged at 17,000 x G for 10 minutes at 4°C. Supernatant was removed and frozen at -80°C as the cytosolic fraction. Pellets were resuspended with cell lysis buffer, briefly sonicated on ice with a probe tip sonicator, and frozen at -80°C as the crude nuclear fraction.

*Lipolysis assays*

Isolated adipocytes were treated with vehicle or 1 μM AS-1842856 for 4 hours in lipolysis assay buffer (145 mmol/L NaCl, 5.4 mmol/L KCl, 1.4 mmol/L $CaCl_2$, 1.4 mmol/L $MgSO_4$, 0.2 mmol/L $NaH_2PO_4$, 5 mmol/L glucose, 2.5% BSA, 100 nmol/L adenosine, 10 mmol/L HEPES, pH 7.4) in a 37°C water bath with mild agitation. Adipocytes were washed twice with assay buffer without adenosine and separated into assay tubes with ~100,000 cells per assay in a total assay volume of 200 μL. Adipocytes were treated with vehicle or 100 nmol/L CCPA followed by the addition of 1

U/mL adenosine deaminase to all lipolysis assays and incubation at 37°C with mild agitation for 15 minutes. Adipocytes were then treated with vehicle or 30 nmol/L isoproterenol and incubated for an additional 30 minutes. Assays were centrifuged for 1 minute at 200 x *g*, adipocytes were aspirated off, and assay buffer was collected and assessed for NEFA concentration. All assays were performed in triplicate, and each N represents independently isolated adipocytes.

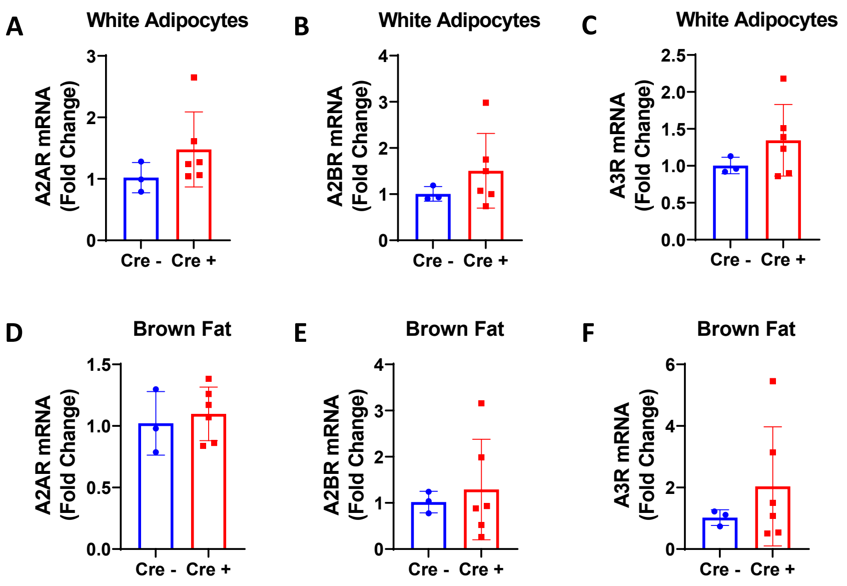


**Supplemental Figure 1. Knockout of A1R does not induce compensatory changes in other adenosine receptors.** (A-C) mRNA expression of adenosine receptors A2A (A), A2B (B), and A3 (C) in isolated gonadal adipocytes. (D-F) mRNA expression of adenosine receptors A2A (D), A2B (E), and A3 (F) in brown adipose tissue.

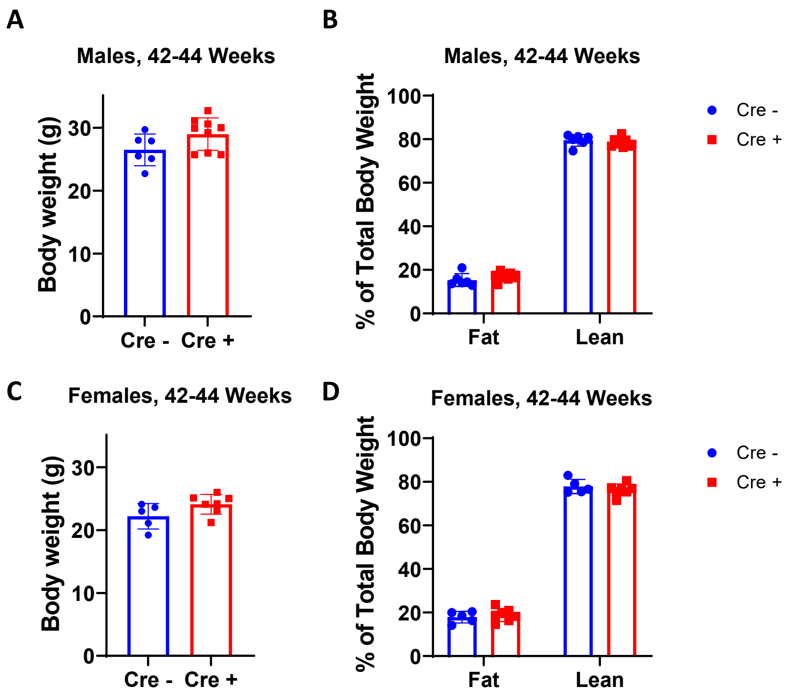


**Supplemental Figure 2. Inducible knockout of adipose A1R does not affect body weight or body mass distribution in aged mice.** (A-D) Cre-negative WT (blue) and cre-positive FAdora1-/- (red) mice were administered tamoxifen at 8 weeks of age. At approximately 42-44 weeks of age, body weight was measured for male (A) and female (C) mice, and body mass distribution was measured by EchoMRI for male (B) and female (D) mice.

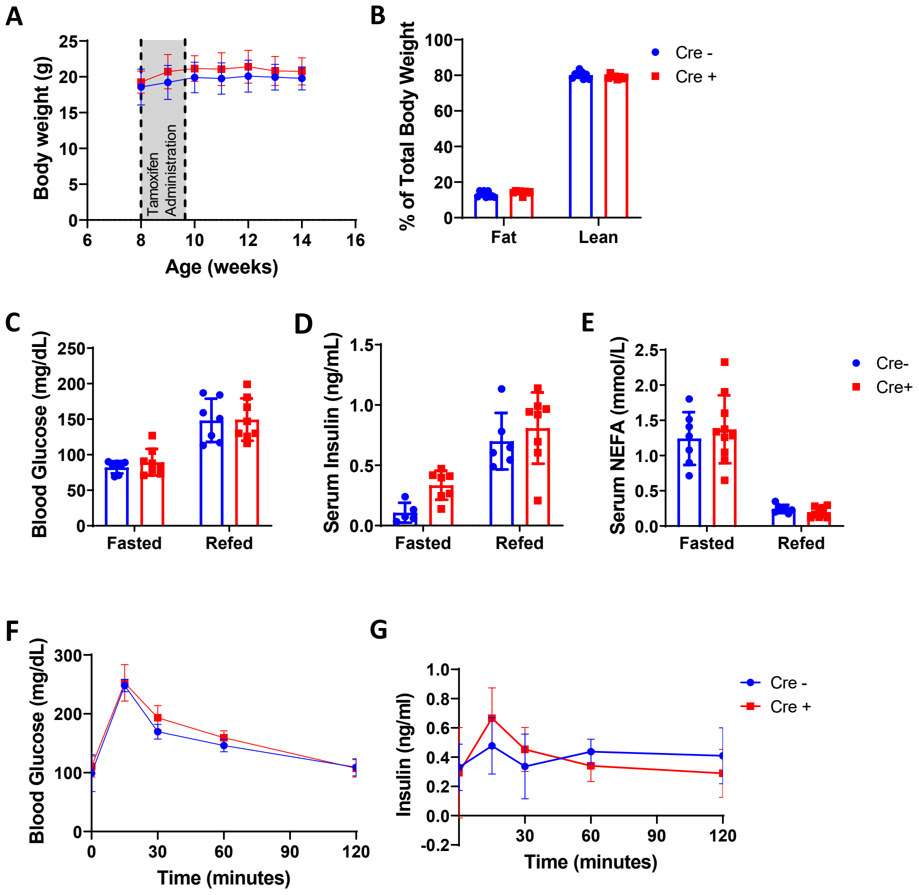


**Supplemental figure 3. Female chow fed FAdora1-/- mice do not display alterations in basal metabolic parameters.** (A) Body weight of female cre-negative WT (blue) and cre-positive FAdora1-/- mice (red) starting at the time of tamoxifen administration at 8 weeks. Gray box indicates timing of tamoxifen. (B) Fat and lean mass distribution in male WT and FAdora1-/- mice 6 weeks after the start of tamoxifen administration measured by EchoMRI. (C-E) Blood glucose (C), serum insulin (D), and serum NEFA (E) levels in female WT and FAdora1-/- mice after an overnight fast (fasted) and 4-hour refeeding period (refed). (F, G) Blood glucose (F) and serum insulin (G) in female WT and FAdora1-/- mice during an i.p. glucose tolerance test with 1 g/kg glucose. Error bars represent SD.

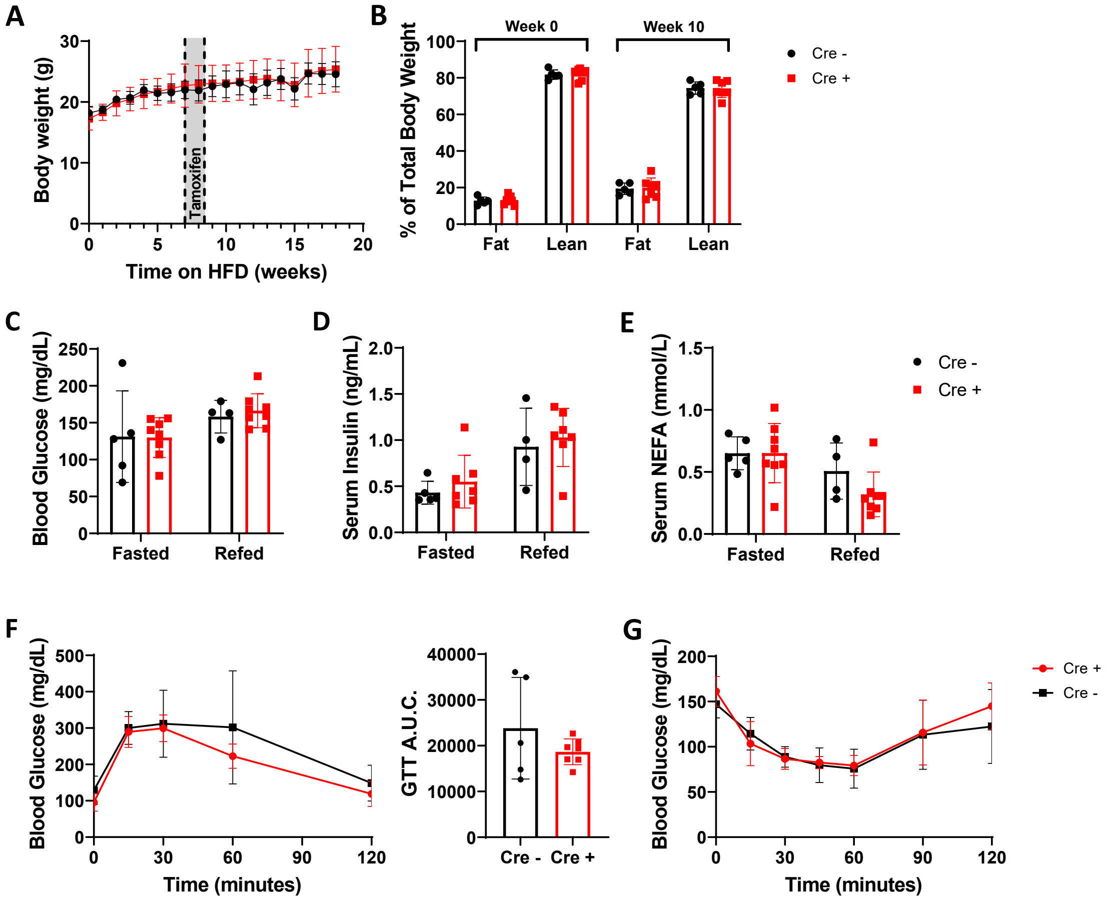


**Supplemental figure 4. Female high-fat diet fed Fadora1-/- mice are not different from WT mice in basal metabolic parameters.** (A) Body weight of female cre-negative WT (black) and cre-positive FAdora1-/- mice (red) starting at the beginning of high-fat diet feeding at 8 weeks. Gray box indicates timing of tamoxifen administration. (B) Fat and lean mass distribution in female WT and FAdora1-/- mice before and after 10 weeks of high-fat diet feeding measured by EchoMRI. (C-G) Data were collected for female WT and FAdora1-/- mice after 12-15 weeks of high-fat diet feeding. (C-E) Blood glucose (C), serum insulin (D), and serum NEFA (E) following an overnight fast (fasted) and 4-hour feeding period (refed). (F) Blood glucose and area under the curve during an i.p. glucose tolerance test with 1 g/kg glucose. (G) Blood glucose during an i.p. insulin tolerance test with 0.75 IU/kg insulin. Error bars represent SD.

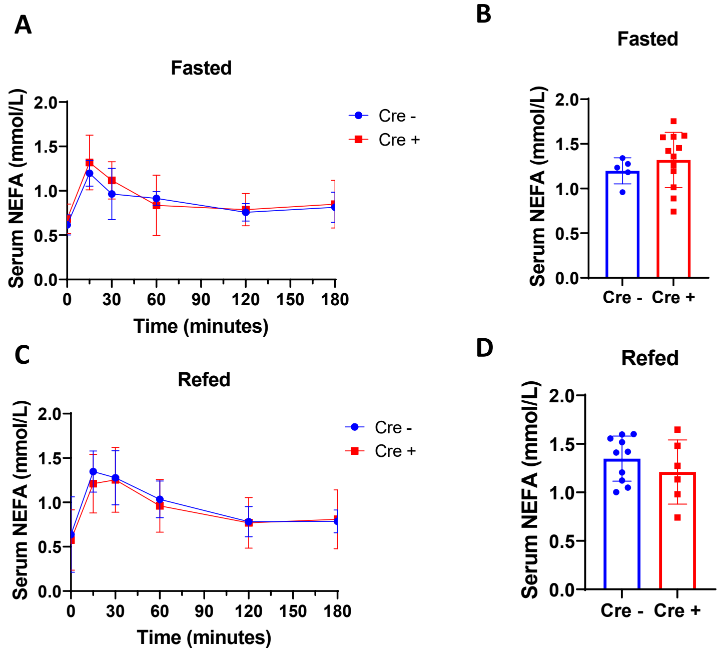


**Supplemental figure 5. Female FAdora1-/- do not display differences in lipolysis under fasted or refed conditions.** (A,B) Serum NEFA in 6-hour fasted female cre-negative WT (blue) and cre-positive FAdora1-/- (red) mice at the indicated time points after i.p. injection of 10 mg/kg isoproterenol (A) or at 15 minutes after injection with isoproterenol (B). (C,D) Same as in (A,B) for mice that were fasted overnight and refed for 4 hours.
